# Primary bovine embryonic fibroblasts support seasonal influenza A virus infection and demonstrate variable fitness of HPAI H5N1

**DOI:** 10.1101/2025.07.26.666677

**Authors:** Grace K. Wenger, Deann T. Snyder, Justin R. Prigge, Allyson H. Turner, Sara A. Jaffrani, Edward E. Schmidt, Emily A. Bruce, Emma K. Loveday

**Author notes:** Emma K. Loveday, **Email:**. **Author Contributions:** EAB, and EKL designed research; GKW, DS, JRP, AHT, SAJ, EAB, and EKL performed research; EAB and EES contributed new tools and reagents; GKW, DS, JRP, AHT, SAJ, EAB, and EKL analyzed data; and GKW and EKL wrote the paper. All authors contributed to editing and revising the manuscript. **Competing Interest Statement:** The authors declare no competing interests.

## Abstract

The emergence of highly pathogenic avian influenza (HPAI) H5N1 (clade 2.3.4.4b, genotype B3.13) in dairy cattle presents substantial challenges to the agricultural sector and public health. Mechanistic studies of infection and transmission in cattle have proven difficult due to animal handling restrictions as well as limited availability of established cell culture models. Primary <u>B</u>ovin<u>e E</u>mbryonic <u>F</u>ibroblasts (BeEFs) were collected from a Montana cow and are investigated here as a model to study influenza A virus (IAV) infection dynamics. We compared sialylation profiles, infectious virus production, viral replication, and plaque morphology in both BeEFs and chicken DF-1 cells following infection with the bovine HPAI H5N1 and an earlier 2.3.4.4b genotype (B1.1) isolated in 2022. The data presented here show increased viral fitness of the bovine origin HPAI H5N1 strains across multiple species and bovine susceptibility to human seasonal IAV. This study highlights the ability of BeEFs to serve as a model for studying IAV infections in bovine hosts.

## Introduction

Since March 2024, highly pathogenic avian influenza (HPAI) H5N1 (clade 2.3.4.4b) has been identified in 1,048 U.S. dairy cattle herds across 17 states, resulting in reduced milk production, decreased quality, and mastitis among infected animals (1, 2). This widespread transmission poses serious economic threats to agriculture, as control measures and necessary biosecurity enhancements would cause considerable financial burdens on dairy operations and related industries (3, 4). Furthermore, infected mammary glands release high concentrations of virus into milk, posing a significant risk of infection to humans exposed to unpasteurized milk (1). Since the emergence of the virus in cattle, dozens of human cases stemming from exposure to infected cattle have been documented (1, 2, 5, 6). However, in depth studies of HPAI H5N1 within bovine hosts is difficult due to animal handling restrictions and a lack of well characterized primary cell culture models. This scarcity is partly due to the recent emergence of influenza A viruses (IAV) in cattle, which has left the field with few established in vitro models. Nonetheless, these studies are critical for advancing our understanding of the molecular mechanisms, viral tropism, and evolutionary potential of IAV in bovine hosts.

To improve understanding of HPAI H5N1 infection dynamics in cattle, we acquired <u>B</u>ovin<u>e E</u>mbryonic <u>F</u>ibroblasts (BeEFs) from a cow of unknown breed in Montana and determined their susceptibility and permissivity compared to DF-1s, a chicken fibroblast cell line, infected with the same HPAI H5N1 viruses from multiple clades. We investigated the presence of sialic acid receptors associated with avian- and mammalian-origin IAV infection, identifying α-2,3 receptors required for avian-type influenza infection on both cell types and limited recognition of α-2,6 receptors in the BeEFs, similar to previously characterized profiles of mammary tissue (7, 8). We compared growth curves using an avian-origin H5N1 isolated before the cattle outbreak, A/Bald Eagle/Florida/W22-134/2022/H5N1 (genotype B1.1), (BEFL), to a current bovine H5N1 isolate, A/Bovine/Ohio/B24OSU-439/2024/H5N1 (genotype B3.13), (BOOH). We also tested the ability of BeEFs to support infection with human seasonal IAVs and found differential susceptibility between previous and recently circulating human seasonal strains. Overall, our results demonstrate that the newly circulating bovine H5N1 strains have increased fitness in both bovine and chicken models of infection. Furthermore, our data shows that bovine cells are susceptible and permissive to recently circulating human seasonal influenza A strains, demonstrating the potential capacity for cattle to operate as mixing vessels for influenza A strains.

## Results and Discussion

### Characterizing BeEF and DF-1 cells following HPAI H5N1 infection

To determine if BeEFs are susceptible to HPAI H5N1 and can function as a potential model system for studying HPAI infection dynamics in cattle, we performed a comparative study of BeEF and DF-1 cells infected with HPAI H5N1 strains isolated before (BEFL) and during (BOOH) the HPAI bovine outbreak (**Figure 1A**). Sialylation profiling was performed on both cell types followed by growth curves to determine the efficiency and fitness of HPAI H5N1 across species.

**Figure 1:**
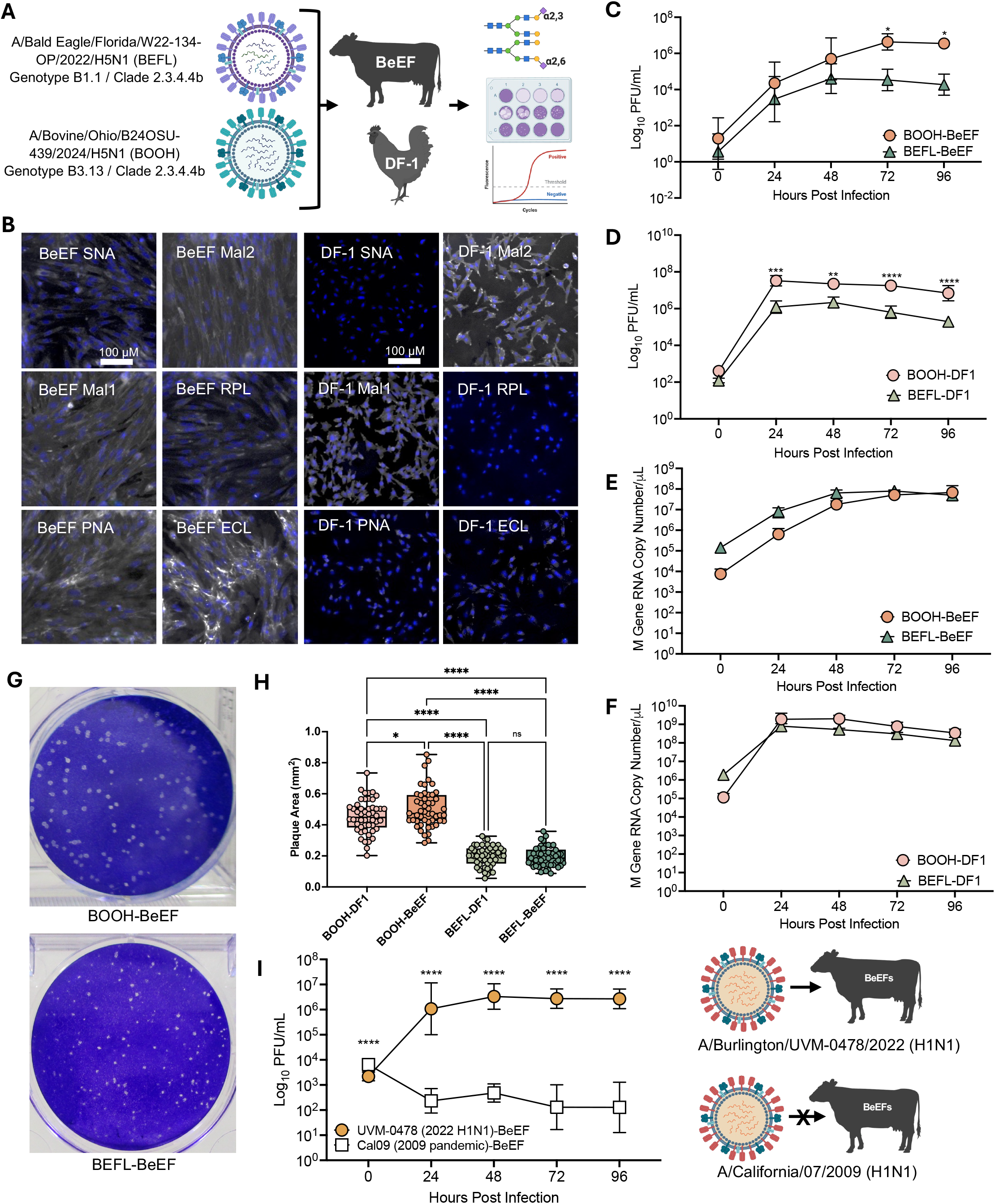
Characterization of HPAI H5N1 viral growth in BeEFs and DF-1 cells. **(A)** A schematic of the experimental design: Infection of BeEF and DF-1 cells with H5N1 isolated before (BEFL) and during (BOOH) the bovine HPAI outbreak. Cells were analyzed for presence of different sialic acid receptors, infectious virus production, and viral replication **(B)** Sialylation profiling displaying lectin-binding (white) of α-2,3 and α-2,6 specific lectins to BeEF and DF-1 cells. Representative images of each lectin are shown at the same magnification. Scale bar = 100 μm. Growth curves (MOI 0.1) of BOOH and BEFL on BeEFS **(C)** and DF-1s **(D)** show increased fitness of BOOH in both cell types. Data represent a minimum of two independent replicates analyzed by two-way ANOVA. Viral replication as measured by RT-qPCR for the viral M gene in total RNA isolated from BeEFs **(E)** and DF-1s **(F)**. Data represent a minimum of two independent replicates analyzed by two-way ANOVA. **(G)** Representative images of plaques formed by BOOH and BEFL infection of BeEFs. **(H)** Comparison of plaque areas reveal significantly reduced plaque sizes for BEFL in both BeEFs and DF-1 compared to BOOH. A minimum of 50 plaques were analyzed for each virus and analyzed by one-way ANOVA. **(I)** Growth curve of Cal09 H1N1 and UVM-0478 H1N1 on BeEFs. Data represent a minimum of three independent replicates analyzed by two-way ANOVA. * p < 0.05, ** p < 0.01, ***p < 0.001 ****p < 0.0001.

### Avian-origin IAV receptors are present on BeEF cells

Characterizing receptor prevalence across different cell types provides critical insight into IAV infection dynamics in multiple species. Current studies show that cow mammary glands exhibit mixed sialic acid profiles, expressing predominantly α-2,3 sialic acid receptors with a lower abundance of α-2,6 receptors (7–9). We compared fluorescence images of BeEF and DF-1 cells exposed to α-2,6-(SNA) and α-2,3-(MAL1, MAL2, and RPL) specific lectins (**Figure 1B**). BeEFs displayed clear binding to MAL1 and MAL2 but also had faint binding to SNA. BeEFs also displayed high levels of binding to PNA and ECL, which target O-glycans and β-1,4 Galactose, respectively, that lack terminal sialic acids. Similarly, DF-1 cells demonstrated clear binding to MAL1 and MAL2 lectins but had limited binding to SNA. In contrast to BeEFs, binding of PNA and ECL to DF1 cells was minimal. These results highlight the strong presence of α-2,3 linkages in the BeEFs, indicating a similar sialylation profile to previously characterized mammary tissue in dairy cattle (7–9).

### Bovine-origin H5N1 shows increased fitness in bovine and chicken primary cells

To assess potential differences in viral replication between the BEFL and BOOH strains, growth curves were performed following low multiplicity of infection (MOI) of the BeEFs (**Figure 1C**). Growth curves revealed a reduction in BEFL virus production at multiple time points compared to BOOH. Growth rates were also analyzed in DF-1 cells, a cell type in which bovine-origin H5N1 has been previously characterized (10), with similar results, although both the BEFL and BOOH strains produced higher titers overall in the DF-1 cells (**Figure 1D**).

Quantification of viral M gene in total RNA from infected cells revealed efficient viral infection and replication with no significant difference in BeEF (**Figure 1E**) or DF-1 cells (**Figure 1F**). To further evaluate fitness differences following observations made during the viral growth curves, we analyzed plaque sizes from virus collected from BeEFs infected with BOOH and BEFL and titered on MST cells, with representative images shown in **Figure 1G**. Plaques from BeEF and DF-1 cells infected with BOOH were significantly larger than plaques from BEFL infected cells, demonstrating increased fitness of the BOOH H5N1 strain across multiple species (**Figure 1H**). Furthermore, the plaques produced from BOOH infected BeEFs were significantly larger than those from BOOH infected DF-1 cells further highlighting the increased viral fitness of BOOH in our bovine cell model. These findings underscore the ongoing zoonotic threat posed by HPAI H5N1 2.3.4.4b strains, whose continued adaptation in avian and mammalian hosts highlights the significant economic risk to the agricultural sector and a growing potential for human spillover and pandemic emergence (1).

### Bovine primary cells are susceptible to recently circulating human seasonal IAV

Previous infections of bovine mammary cells with the IAV PR8 strain found that virus could be secreted in the milk for multiple days (11). In addition, seropositivity and isolation of human and human-like IAVs from natural infection in cattle have previously been reported (12). Therefore, we investigated the susceptibility of BeEFs to human seasonal IAV strains. The BeEFs were infected with the 2009 pandemic strain, A/California/07/2009 (Cal09) H1N1 and a recent seasonal strain isolated from an IAV positive clinical specimen, A/Burlington/UVM-0478/2022 (UVM-0478) H1N1 (13). Viral replication was measured via plaque assays following low-MOI growth curves (**Figure 1I**). The growth curves revealed that BeEFs are not susceptible to Cal09 H1N1 with limited detectable plaques over 96 hours post infection (hpi), most likely representing remaining inoculum. Notably, BeEFs were fully susceptible and permissive to UVM-0478. We observed a multi-log increase in infectious virus released at 24 hpi that was maintained over 96 hpi. (**Figure 1I**). Evidence of BeEF cell susceptibility to currently circulating human seasonal IAVs as well as H5N1 HPAI demonstrates the potential for simultaneous infection and reassortment of these strains.

### Conclusion: BeEFs as a potential model for studying HPAI H5N1 infection dynamics in cattle

The emergence of HPAI H5N1 2.3.4.4b strains across a variety of hosts poses a significant threat to both the agricultural industry and human health. While zoonotic spillover events into humans are rare due to restrictions that limit viral replication, human infection with bovine origin H5N1 has occurred multiple times over the past year (5, 6). Continued adaptation and evolution of HPAI H5N1 2.3.4.4b strains within avian, bovine, and human hosts indicates the virus’s continued zoonotic potential, increasing pandemic potential. However, there is a lack of well described *in-vitro* bovine models to study IAV. To address this need, we compared HPAI H5N1 infection dynamics in primary BeEFs to chicken DF-1 cells to determine their capacity to support infection with bovine-origin and avian-origin HPAI H5N1. Similar to recent studies in bovine mammary cells, the BeEFs support higher replication of bovine-origin HPAI strains compared to earlier clade avian-origin viruses (9). Furthermore, we have demonstrated sustained infection of recently circulating (2022) human-origin IAV in the BeEFs, although they remain refractory to infection with the 2009 pandemic H1N1 strain. Previous research demonstrated slight permissivity of bovine mammary cells to a seasonal H3N2 virus isolated in 2013 suggesting that evolution of human IAV strains may have expanded their capacity to infect bovine hosts (9). In addition, we found sialic acid expression in the BeEFs was consistent with previous reports on bovine mammary tissue further validating BeEFs as a model for studying influenza A virus within a bovine host. Taken together, our findings underscore the value of BeEFs as a biologically relevant in vitro system for studying host-specific adaptations of influenza A viruses in cattle and advancing our understanding of cross-species transmission dynamics.

## Materials and Methods

### Cells lines and maintenance

Madin-Darby canine kidney (MDCK) cells were obtained from Val Le Sage at the University of Pittsburgh and MDCK-SIAT1-TMPRSS2 (MST) cells were obtained from Jesse Bloom at the Fred Hutch Cancer Center (14). Chicken embryonic fibroblast (DF-1) cells were obtained from Steve Baker at Lovelace Biomedical. Bovine embryonic fibroblasts (BeEFs) were provided by Ed Schmidt at Montana State University. All cells grown were maintained in DMEM (Corning) supplemented with 10% FBS (Gibco), 1X penicillin-streptomycin (Gibco), and with (MSU)/without (UVM) 2.5 μg/ml plasmocin (Invivogen). All cells were passed weekly or as needed.

### Harvesting BeEFs

Bovine fetal harvest procedures were reviewed and approved by the MSU IACUC, protocol # 1999-13. To favor high heterozygosity, an “outbred Montana free-range heifer” was purchased at public auction (Bozeman, Montana, Autumn, 1999), verified clear, cycled, and artificially inseminated with bovine semen from a local generic undocumented source. At 45 days post insemination (23 February 2020), a single early fetus in intact decidua was harvested by manual palpation into a liter of ice-cold PBS containing 2X antibiotic cocktail (“PBS-ab”, Gibco # 15240096). The full decidua was transported to the laboratory in PBS-ab on ice, transferred to a new beaker of PBS-ab, and the maternal decidual tissues were carefully dissected away down to the fetus-containing amniotic sac, which was maintained unbreached. This was surface-disinfected by 70% ethanol-dip for 30 sec and then dipped into a fresh beaker with sterile PBS-ab (to remove ethanol) in a biosafety cabinet and transferred into a sterile culture dish. The amniotic sac was breached and the fetus (∼ 2 cm long) was transferred to a fresh dish. The dermis was gently collected and transferred into a 50 ml tube containing 10 ml 2X trypsin solution (Gibco # 15090046). After 20 min 37°C, 35 ml of DMEM + 10% newborn calf serum (NCS) + 1X antibiotics was added to quench the trypsin. Cells were sedimented by centrifugation (300 x g 10 min), supernatant was aspirated, and cells were dispersed by gentle trituration with 20 ml of DMEM + 10% NCS + 1X antibiotics and plated onto 10 × 10-cm cell culture dishes in the same medium. Media was replaced the next day. One day later, the cells had reached near-confluence; they were collected with trypsin, frozen as “P0” stocks, and maintained in liquid nitrogen thereafter. Diploid bovine karyotype (n = 60 chr) was verified.

### Influenza strains and production

The following strains of avian influenza viruses were obtained from Richard Webby (St Jude’s, Memphis, TN): A/bovine/Ohio/B24OSU-439/2024 H5N1 and A/bald eagle/Florida/W22-134-OP/2022 H5N1. The following influenza virus was obtained from Jess Crothers at the University of Vermont (Burlington, VT): A/Burlington/UVM-0478/2022 (UVM-0478) H1N1. A/California/07/2009 (Cal09) H1N1 was obtained from Chris Brooke at the University of Illinois at Urbana-Champaign. HPAI H5N1 viral stocks were generated from MDCK cells in infection media containing DMEM (Corning) supplemented with 0.3% BSA (MP Biomedicals), 1X penicillin-streptomycin (Gibco), 1mM HEPES (Gibco). For the H1N1 viruses 2ug/TPCK-trypsin (Worthington Biomedical) was also added to the infection media. Supernatants were collected and centrifuged at 3,000 g for 5 min to remove cellular debris. The clarified viral supernatant was titered by plaque assay on MST cells, sequence verified and then used for all subsequent infections. All experiments with H5N1 strains were performed in the Jutila Research Biosafety Level 3 Lab at Montana State University, Bozeman, Montana. Seasonal IAV stocks were produced as previously described (13). Briefly, viral stocks were grown on MSTs in a T150 flask (Corning) and infected at an MOI of 0.001 in OptiMEM (Gibco) plus 1 ug/ml TPCK trypsin (Sigma-Aldrich) and incubated at 37°C until ∼50% CPE was observed.

### BeEF and DF-1 cell infections

Cells were counted the day of infection followed by inoculation at an MOI of 0.1. TPCK-Trypsin was added to infection media at 1 μg/ml for H1N1 infections but omitted for H5N1 strains. Cells were washed twice with PBS then inoculated for one hour at 37C. Following infection, inoculum was removed and replaced with fresh infection media. To collect the 0 hour time point, the fresh media was added for 10 minutes prior to collecting. The cells were returned to 37C and supernatants were collected from individual wells every 24 hours for the duration of the experiment (96 hours).

### Quantification of viral titers by plaque assay

HPAI H5N1 titers were determined via plaque assays with MST cells, as previously described (15). Briefly, MST cells were cultured in complete DMEM, seeded onto 6-well plates at 7.5e5 cells per well and grown to confluence overnight. They were then incubated with serially diluted viral inoculum in 200 μl infection media for 1 h. Following inoculation, cells were over-layered with 1.5% carboxymethylcellulose, DMEM, 0.2mg/ml DEAE-dextran, 0.3% BSA, 1X penicillin-streptomycin, 1mM HEPES and 3.7g/L sodium bicarbonate and incubated for 3–4 days. Cells were then fixed with 4% paraformaldehyde for 30 minutes and stained with 0.1% crystal violet in water for 15-20 minutes. Plaques were counted and the titer was calculated based on the volume of inoculum plated and dilution counted.

Seasonal H1N1 titers were performed as previously described (13). Briefly, MST cells were seeded at 1e6 cells per well and inoculated with serial ten-fold dilutions of each sample in DMEM. After one hour, cells were overlayed with 2 ml of a warmed 1:1 mixture of 2.4% Avicel RC-591 NF (Dupont) + 1X DMEM (Corning) + 1 μg/ml TPCK trypsin (Sigma-Aldrich). Cells were returned to 37C for 48 hours, and care was taken not to disturb the plates during this period. Finally, cells were washed with PBS (Corning), fixed with 4% formaldehyde (Honeywell) for 20 minutes and stained with 0.1% crystal violet (Fisher) for five minutes. Plates were rinsed three times with water and allowed to dry before plaques were counted to determine viral titer.

Replicate titer values at each time point and condition were compiled and used to generate viral growth curves. Statistical comparisons of viral replication across timepoints and conditions were analyzed with a two-way ANOVA. Significance was defined as p < 0.05. Statistical analyses were completed using Prism.

### M gene abundance in total RNA by RT-qPCR

RNA was extracted from cell lysates using the Invitrogen PureLink RNA mini kit (Invitrogen). RNA copy number of the IAV matrix protein gene (M gene) was determined using a standard curve and TaqMan RT-qPCR assay as previously described (15, 16). The amplification primer sequence was as follows: M gene forward, 59-GACCRATCCTGTCACCTCTGAC-39, and M gene reverse, 59-AGGGCATTCTGGACAAATCGTCTA-39. The sequence of the M gene TaqMan probe was 59-/FAM/TGCAGTCCTCGCTCACTGGGCACG/BHQ1/-39. Working stocks of the primers and probe (Eurofins Operon) were prepared at 25 mM and 10mM, respectively, for use in the RT-qPCR. Samples were amplified using a SuperScript III Platinum one-step RT-qPCR kit (Invitrogen) with a final reaction volume of 25 mL. Each reaction mix contained 400 nM M gene forward and reverse primers, 200 nM M gene TaqMan probe, 0.05 mM ROX reference dye, 5U/mL Superase RNase inhibitor (Invitrogen), and 5 μL of RNA template. Thermocycling was performed in an RT-qPCR machine (QuantStudio 3; Applied Biosystems) with the following cycling conditions: 1 cycle for 30 min at 60°C, 1 cycle for 2 min at 95°C, and 40 cycles between 15 s at 95°C and 1 min at 60°C.

### Plaque Size Quantification

Images of plaques from A/Bovine/Ohio/B24OSU-439/2024/H5N1 (BOOH) and A/Bald Eagle/Florida/W22-134/2022/H5N1 (BEFL) infections of BeEF and DF-1 cells were captured following infection and staining on a light tracing box with built in scale (Amazon). Images were imported to Fiji (Image J) and the appropriate scale was calibrated using the included scale bar. For each virus-cell type combination, fifty distinct plaques were outlined manually and measured using the “Analyze Particle” function. The resulting measurements were compiled and analyzed using a one-way ANOVA. Significance was defined as p < 0.05. Statistical analyses were completed using Prism.

### Sialylation Profiling by Lectin Staining

Chicken embryonic fibroblasts (DF-1) and bovine embryonic fibroblasts (BeEF) were seeded at 1e5 cells per well on an 8-well LabTek chambered coverslips (Thermo Fisher Scientific) and incubated for 24 hours until adherent. Cells were washed with PBS and then fixed using 4% PFA for 30 minutes at room temperature. To reduce non-specific binding, the cells were washed twice with PBS and incubated in a blocking solution of 2% BSA in PBS for 10 minutes at room temperature.

Lectin staining was performed using the Alpha 2,3/2,6 Sialylation Profiling Kit (ZBiotech) with modifications to adapt the manufacturer’s protocol to fixed adherent cells. Each of 6 wells per cell type were stained with a distinct biotinylated lectin (100 μl) at a final concentration of 0.125 μg/100 μL in PBS and incubated at room temperature for 1 hour. After primary lectin incubation, cells were washed several times with PBS and then incubated with FITC-conjugated streptavidin (100 μl) at 7 μg/mL in PBS at room temperature for 1 hour. Cells were washed several times with PBS and then nuclei were stained with Hoescht solution (100 μl) at 2 μg/mL in PBS and incubated at room temperature for 10 minutes. Cells were washed several times and slides were stored with PBS until imaging.

Fluorescence microscopy was conducted using a 20X objective lens. Lookup table (LUT) settings were normalized across all images to enable direct comparison of the images. All staining experiments were repeated using independently plated cells with representative images shown. Additional images were acquired with adjusted LUT settings.

## Acknowledgments

This work was supported by grants from the National Institutes of Health (NIH) under NIAID from the Johns Hopkins Centers of Excellence in Influenza Research and Research (CEIRR) Option 13E HHS N7593021C00045 awarded to EKL, NIAID (R21 AI178432-01) awarded to EKL, and NIGMS (P20GM125498) awarded to EKL and EAB. The authors acknowledge critical roles of James Thompson (MSU) for animal acquisition and management, and of Dr. Bruce Sorensen (Sorensen Large Animal Veterinary Hospital, Belgrade, Montana) for artificial insemination and embryologic procedures.

Development of the BeEF cell line was further supported by grants from the US Department of Agriculture (#35208-11567) and the Montana Agricultural Experiment Station (#MONB00443) to EES. We thank Richard Webby at St. Jude Children’s Research Hospital for the bald eagle and bovine H5N1 influenza strains, Chris Brooke at the University of Illinois at Urbana-Champaign for the California 2009 H1N1 influenza strain and Jessica Crothers at the University of Vermont for providing the 2022 H1N1 clinical specimen. We thank Valerie Le Sage at the University of Pittsburgh for MDCK cells, Jesse Bloom at the Fred Hutch Cancer Center for the MDCK-SIAT1-TMPRSS2, and Steve Baker at Lovelace Biomedical for the DF-1 cells.

